# A receptor-like kinase controls plasmodesmal transport of conserved 30K viral movement proteins through phosphorylation

**DOI:** 10.64898/2026.02.10.705008

**Authors:** Juan Martínez-Sáez, Ángel Chávez, Andrea Restrepo-Escobar, T. Moritz Schladt, Wolf B. Frommer, Waltraud X. Schulze, Manuel Miras

## Abstract

Intercellular viral movement in plants is mediated by movement proteins (MPs) that modulate plasmodesmata (PD) enabling cell-to-cell and systemic trafficking. Although phosphorylation has long been implicated in the regulation of MP localization and activity, the identity of host kinases and the interface with immune signaling have remain unresolved. Here, we identified the *Arabidopsis thaliana* lectin receptor-like kinase RDA2 as a PD-associated regulatory component of viral movement. Using proximity labeling, we detected RDA2 as a proximal interactor of the tobacco mosaic virus (TMV) MP, and show that RDA2 directly phosphorylates MP at multiple sites *in vitro*. Phosphorylation at threonine 75 is required for efficient PD targeting and intercellular movement, while phospho-dead mutants failed to complement viral spread. Loss of RDA2 enhanced MP mobility and increased TMV accumulation *in planta*, indicating that RDA2 modulates PD transport during the infection. RDA2 also interacted with and phosphorylated the movement protein of cucumber mosaic virus, implicating that this regulatory mechanism extends across members of the 30K MP superfamily. Our findings demonstrate that a plasma membrane receptor-like kinase can directly modify viral movement proteins, establishing a mechanistic link between receptor-mediated immune signaling and the post-translational control of symplasmic connectivity.

**One-sentence summary:** RDA2, a lectin receptor-like kinase, directly phosphorylates conserved 30K viral movement proteins to control their plasmodesmal targeting and restrict cell-to-cell viral spread in plants

## Introduction

Intercellular communication is essential for plant growth, development, and immunity ^1–4^. In higher plants, this communication relies largely on plasmodesmata (PD)—specialized membrane-lined channels that traverse cell walls and establish cytoplasmic continuity between adjacent cells ^5^. Through PD, plants exchange metabolites, proteins, RNAs, and diverse signaling molecules, ensuring coordination of physiological processes at the whole-organism level ^1,6–10^. However, the same symplasmic connectivity that supports plant development also provides a route for pathogen invasion ^11–13^.

Plant viruses exploit PD for cell-to-cell movement. To spread between cells, viruses encode movement proteins (MPs) that modify PD structure and permeability, enabling viral ribonucleoprotein trafficking ^14^. Many of these MPs, including those from tobacco mosaic virus (TMV) and cucumber mosaic virus (CMV), belong to the evolutionarily conserved 30K superfamily ^15^. Members of this family share a conserved jelly-roll structural core, but differ substantially in their host interactions, reflecting functional diversification within this superfamily ^16^.

Post-translational modifications, particularly phosphorylation, play pivotal roles in regulating MP localization, activity, and trafficking ^17–19^. Decades of research have mapped multiple phosphorylation sites on TMV MP, revealing that phosphorylation can act as a molecular switch controlling movement capacity ^17,20–22^. Although the *Arabidopsis thaliana* PD-associated casein kinase I–like enzyme (PAPK1) can phosphorylate C-terminal MP residues *in vitro* ^22^, the host kinases responsible for modifying the regulatory sites that control MP targeting and trafficking *in planta*—and how these phosphorylation events are connected to antiviral signaling—remain unknown.

Receptor-like kinases (RLKs) are compelling candidates for to fulfil this missing regulatory link. RLKs are plasma membrane-anchored receptors that integrate extracellular detection with intracellular phosphorylation cascades to regulate development, stress responses, and PD permeability ^23,24^. The leucine-rich repeat RLK BARELY ANY MERISTEM 1 (BAM1), for example, localizes at PD and promotes the cell-to-cell movement of RNA interference signals, and is targeted by viral proteins such as the C4 protein of tomato yellow leaf curl virus (TYLCV) and the silencing suppressor P19 ^25,26^. In addition, BAM1 physically interacts with TMV MP and facilitates early TMV spread in Arabidopsis and *Nicotiana benthamiana* ^27,28^. Yet, despite these examples, no RLK has been shown to directly phosphorylate a viral MP or to control its phosphorylation status.

To uncover host kinases mediating these processes, we applied *in planta* proximity labeling to identify proteins spatially associated with TMV MP. Here, we identify the *A. thaliana* lectin RLK RDA2 (RESISTANT TO DFPM-INHIBITION OF ABSCISIC ACID SIGNALING 2) as a host kinase that integrates immune perception with direct control of viral movement. RDA2, known as a pattern-recognition receptor for microbe-derived sphingoid bases ^29^, localizes to both the plasma membrane (PM) and PD, interacts with MPs from TMV and CMV, and directly phosphorylates them at defined residues. This modification determines the subcellular fate, PD localization, and movement competence of the MPs. Because both MPs belong to the 30K superfamily, our findings reveal a conserved molecular mechanism whereby RLKs link immune receptor signaling to the post-translational control of symplasmic transport, thus providing a new paradigm for virus-host interactions in plants.

## Results

### Deciphering MP-associated host proteins using proximity labeling

To uncover host proteins associated with the TMV MP, we employed proximity labeling using TurboID in Arabidopsis lines expressing TMV MP fused to TurboID and mVenus under the UBIQUITIN10 (UBQ10) promoter. The MP-TurboID-mVenus fusion localized to punctate structures at the cell periphery and overlaying with the PD-associated callose dye (aniline blue), consistent with a localization at PD (**Fig. 1a**). Biotin incubation induced robust biotinylation activity in MP-TurboID-mVenus plants, confirming effective TurboID activity *in vivo* (**Fig. 1b**).

**Fig 1:**
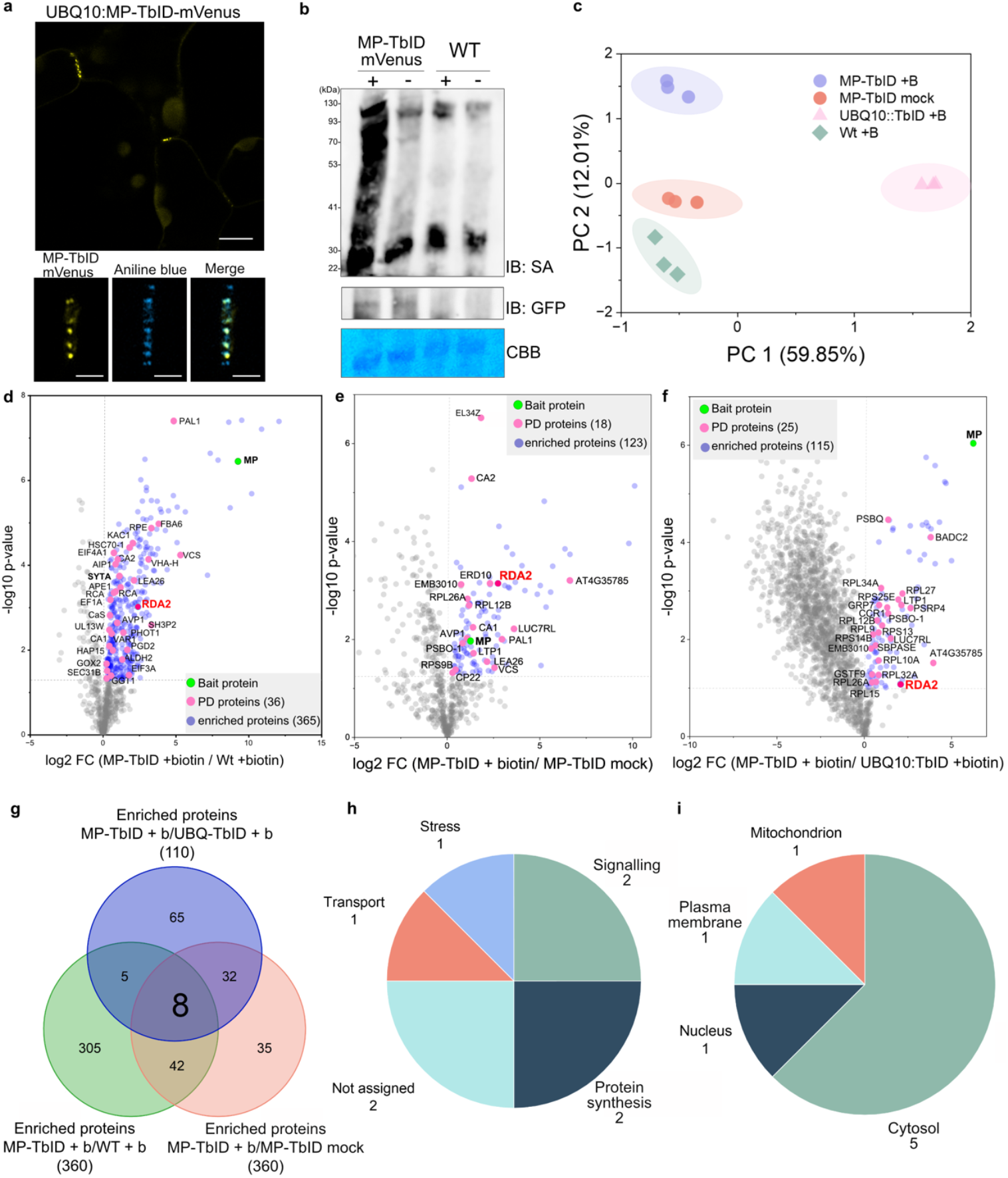
Proximity labeling identified RDA2 among MP-associated proteins *in planta*. **a** Confocal images of *Arabidopsis* UBQ10:MP-TbID-mVenus showing fluorescence at cell boundaries and in punctate structures corresponding to plasmodesmata (PD). Co-staining with aniline blue confirmed colocalization of MP-TbID-mVenus with callose at PD (bottom panels). Scale bars, 20 μm (top) and 10 μm (bottom). **b** Biotinylation activity of MP-TbID-mVenus compared with wild type (WT) controls. Twenty-two-day-old plants were incubated with 50 μM biotin for 1 h prior to extraction. Plant extracts were analyzed by immunoblotting (IB) with streptavidin-HRP (SA) to detect biotinylated proteins and with anti-GFP antibody to confirm expression. Coomassie Brilliant Blue (CBB) staining of the same membrane after transfer shows equal protein loading. **c** Principal component analysis (PCA) of LC-MS/MS proteomic datasets showing clustering of samples according to genotype and biotin treatment. **d-f** Volcano plots showing proteins significantly enriched (p < 0.05) in comparisons between (D) MP-TbID + biotin vs. WT + biotin, (E) MP-TbID + biotin vs. MP-TbID mock, and (F) MP-TbID + biotin vs. UBQ10:TbID + biotin. Viral MP and representative host proteins are highlighted, including the previously reported MP interactor SYTA and the RLK RDA2. **g** Venn diagram showing overlap of enriched proteins identified in the three independent comparisons. Eight proteins were consistently enriched across all datasets, including RDA2. **h** Functional classification of the 8 consistently enriched proteins based on Gene Ontology (GO) annotation. **i** Subcellular localization of the same 8 proteins according to TAIR and SUBA4 annotations.

After optimizing labeling conditions (50 μM biotin for 1 h) (**Supplementray Fig. 1**), we performed streptavidin-based affinity purification followed by LC-MS/MS to identify potential MP interactors. Principal component analysis (PCA) revealed a clear separation between genotypes and between biotin-treated and mock samples (**Fig. 1c**). To obtain a high-confidence candidate list, we applied a strict three-filter approach using different genotypes and biotin-treated versus mock samples. The first filter removed proteins biotinylated non-specifically as a consequence of biotin treatment alone. Comparison of MP-TurboID (biotin) with wild-type (biotin) revealed 361 proteins specifically enriched in MP-TurboID (**Fig. 1d)**. Notably, 35 candidates overlapped with previously reported PD proteomes (PDDB http://pddb.uni-hohenheim.de ^30,31^), including Synaptotagmin A (SYTA), a confirmed TMV MP interactor ^32^ (**Fig. 1e**). The second filter aimed to remove proteins biotinylated as a result of endogenous or low-level TurboID activity, achieved by comparing biotin-treated and mock-treated MP-TurboID samples, identifying 118 enriched proteins, including 17 known PD proteins. Finally, the third filter aimed to exclude highly abundant proteins biotinylated independently of MP by comparing MP-TurboID (biotin) with the UBQ10:TurboID (biotin) line ^33^, which is expressed abundantly in the cytosol, resulting in 111 enriched proteins, 26 of which were previously annotated as PD proteins (**Fig. 1f**). Intersecting the three datasets yielded 9 consistently enriched proteins (**Fig. 1g**), one of which was MP itself, consistent with the known oligomerization of TMV MP ^34,35^. The remaining 8 host candidates included proteins from signaling, stress response, and miscellaneous categories with diverse subcellular localization according to SUBA4 annotation (**Fig. h-i and Supplementary Table 1**).

### Identification of RDA2 as a plasmodesmal receptor-like kinase interacting with TMV MP

Among the eight high-confidence candidates identified by proximity labeling, we focused on RDA2, a lectin receptor-like kinase (LecRLK) with known roles in defense signaling ^29^(**Fig. 1d-f, Fig. 2a**). We first investigated the *in vivo* localization of RDA2 by transiently expressing a C-terminal RDA2-GFP fusion in *N. benthamiana* epidermal cells. As expected for an RLK, the fusion RDA2-GFP localized to the PM (**Fig. 2b**), but closer inspection revealed discrete puncta at the cell periphery (**Fig. 2b**). Quantification using the plasmodesmal enrichment ratio (PD index) confirmed significant PD association (**Fig. 2c**). To further study the association of RDA2 with PD and TMV MP, we co-expressed RDA2-mCherry with TMV MP-GFP. Confocal microscopy showed that both proteins co-localized with aniline-blue-stained PD (**Fig. 2d**). In Arabidopsis *rda2* mutant complemented with UBQ10:RDA2-GFP, RDA2-GFP was similarly detected at the PM and overlaying PD marker (**Supplementary Fig. 2**), supporting that RDA2 localized to PD-associated microdomains *in planta*. The identification of RDA2 as a proximal interactor of TMV MP would indicate a potential regulatory link between viral movement and host receptor-like kinases. Given the established role of LecRLKs in signal transduction, we further investigated whether RDA2 directly interacts with and regulates TMV MP function.

**Figure 2.**
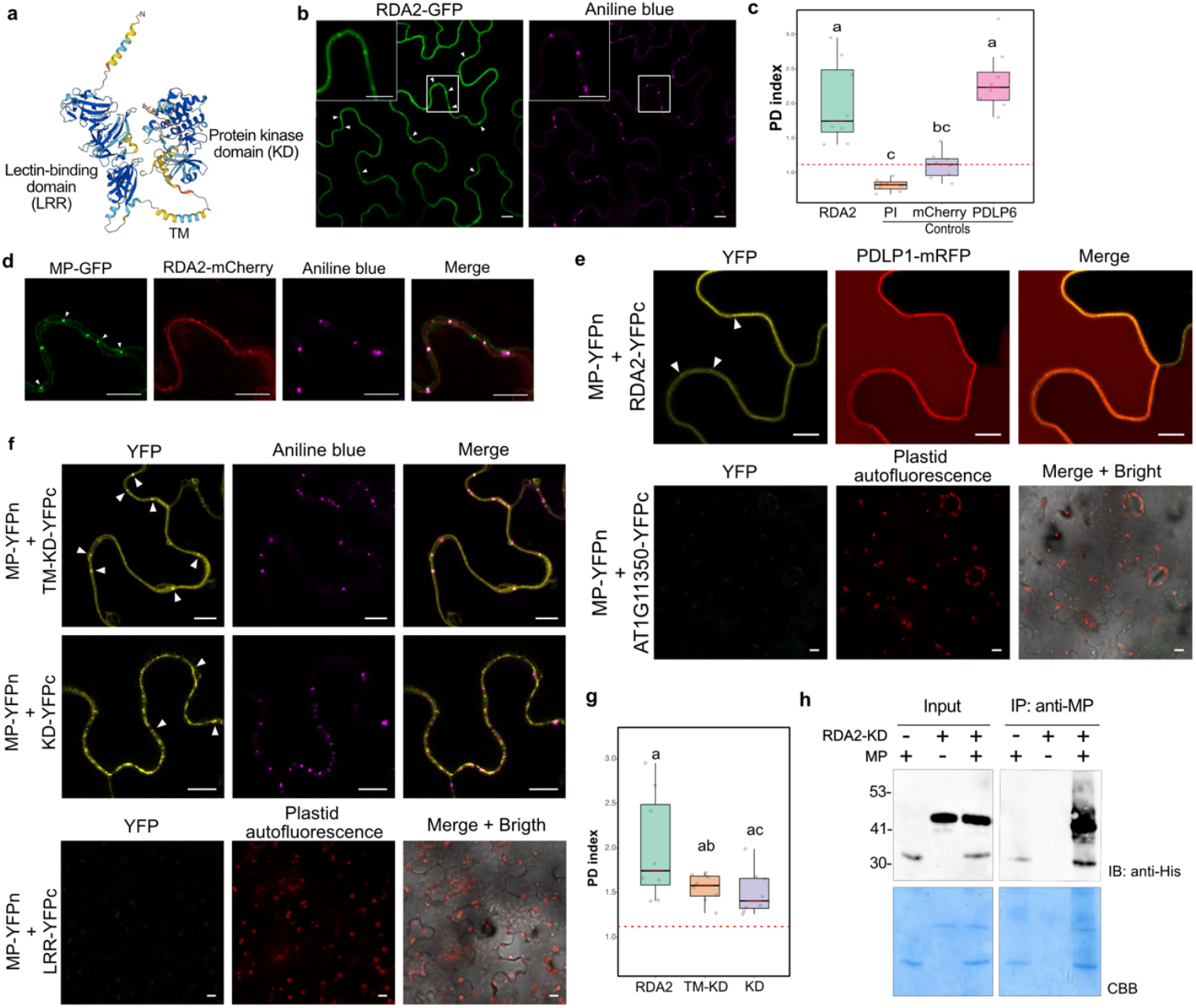
RDA2 localizes to the plasma membrane and plasmodesmata and interacts with TMV movement protein. **a** AlphaFold structural prediction of RDA2 showing lectin-binding (LRR), transmembrane (TM), and kinase (KD) domains. **b** Confocal images of *N. benthamiana* epidermal cells transiently expressing RDA2-GFP. RDA2-GFP fluorescence (green) accumulated at the cell periphery and in discrete puncta overlapping with the PD marker aniline blue (magenta). White arrowheads indicate overlay with aniline blue. Insets show magnified regions. Scale bars: 10 μm. **c** Quantification of PD enrichment (PD index) for RDA2-GFP. Red dashed line indicates the mean PD index of mCherry (cytoplasmic control). PI, propidium iodide (‘membrane impermeant dye’). Boxplots show median and interquartile ranges. Statistical differences were assessed by Kruskal-Wallis test (p < 0.05) followed by Bonferroni post-hoc comparison. Distinct letters indicate statistically significant differences. At least 8 images from three biological replicates were analyzed for each protein. **d** Co-localization of TMV MP-GFP (green) and RDA2-mCherry (red) in *N. benthamiana* epidermal cells. Punctate fluorescence signals (arrowheads) overlapped with aniline-blue-stained PD (magenta). Scale bars: 10 μm. **e** Split YFP revealed interaction of TMV MP (MP-YFPn) with RDA2 (RDA2-YFPc) at PD, marked by PDLP1-mRFP (upper panels). The closest RDA2 homolog, AT1G11350-YFPc, was used as a negative control (lower panels). Scale bars: 10 μm. **f** Split YFP assays using RDA2 truncations showing interaction with TM-KD and KD domains (top, middle), but not with the LRR domain alone (bottom). **g** PD index quantification for each RDA2 fragment. Box plot represents n > 8 cells from three independent experiments. Statistical differences were assessed by Kruskal-Wallis test (p < 0.05) followed by Bonferroni post hoc comparison; different letters indicate statistically significant groups. **h** *In vitro* co-immunoprecipitation confirmed direct interaction between purified His-tagged RDA2-KD and MP, detected by anti-His immunoblot (IB) after pull-down with anti-MP antibody. CBB: same gel stained with Coomassie brilliant blue after transfer. The molecular mass of RDA2-KD-6xHis and TMV MP-6xHis are 42 and 30 kDa, respectively.

To validate whether TMV MP and RDA2 interact, we first employed split-fluorescent protein (split-FP) assays in *N. benthamiana* leaves. MP was fused to the N-terminal half of mVenus (MP-nYFP) and RDA2 to the C-terminal half (RDA2-cYFP). Co-expression of MP-nVenus and RDA2-cVenus resulted in clear YFP fluorescence at the plasma membrane and enriched at PD of epidermal cells, co-localizing with the bona fide PD marker PDLP1 (**Fig. 2e**). In contrast, neither MP-nYFP co-expressed with cYFP alone nor with the closest RDA2 homolog, AT1G11350, produced detectable fluorescence, thereby confirming specificity (**Fig. 2e, Supplementary Fig. 3**). We then examined the MP-RDA2 interaction in more detail. RDA2 is composed of an extracellular lectin domain (LRR) and an intracellular kinase domain (KD) separated by a transmembrane domain (TM) (**Fig. 2a)**. Co-expression of MP-nYFP with KD-cYFP or TM-KD-cYFP both reconstituted YFP fluorescence, whereas the LRR-cYFP fusion did not interact with MP-nYFP. KD alone interacted predominantly in the cytoplasm, whereas TM-KD produced peripheral puncta consistent with PD localization (**Fig. 2f**). Quantification using the PD index confirmed that TM-KD-cYFP showed significantly higher enrichment at PD compared to KD alone or negative controls, and similar to PDLP6 (**Fig. 2g**). These results indicated that RDA2-MP interaction requires the membrane-proximal kinase domain and occurs specifically at PD microdomains, positioning RDA2 as a membrane-anchored receptor sensing MP.

**Figure 3.**
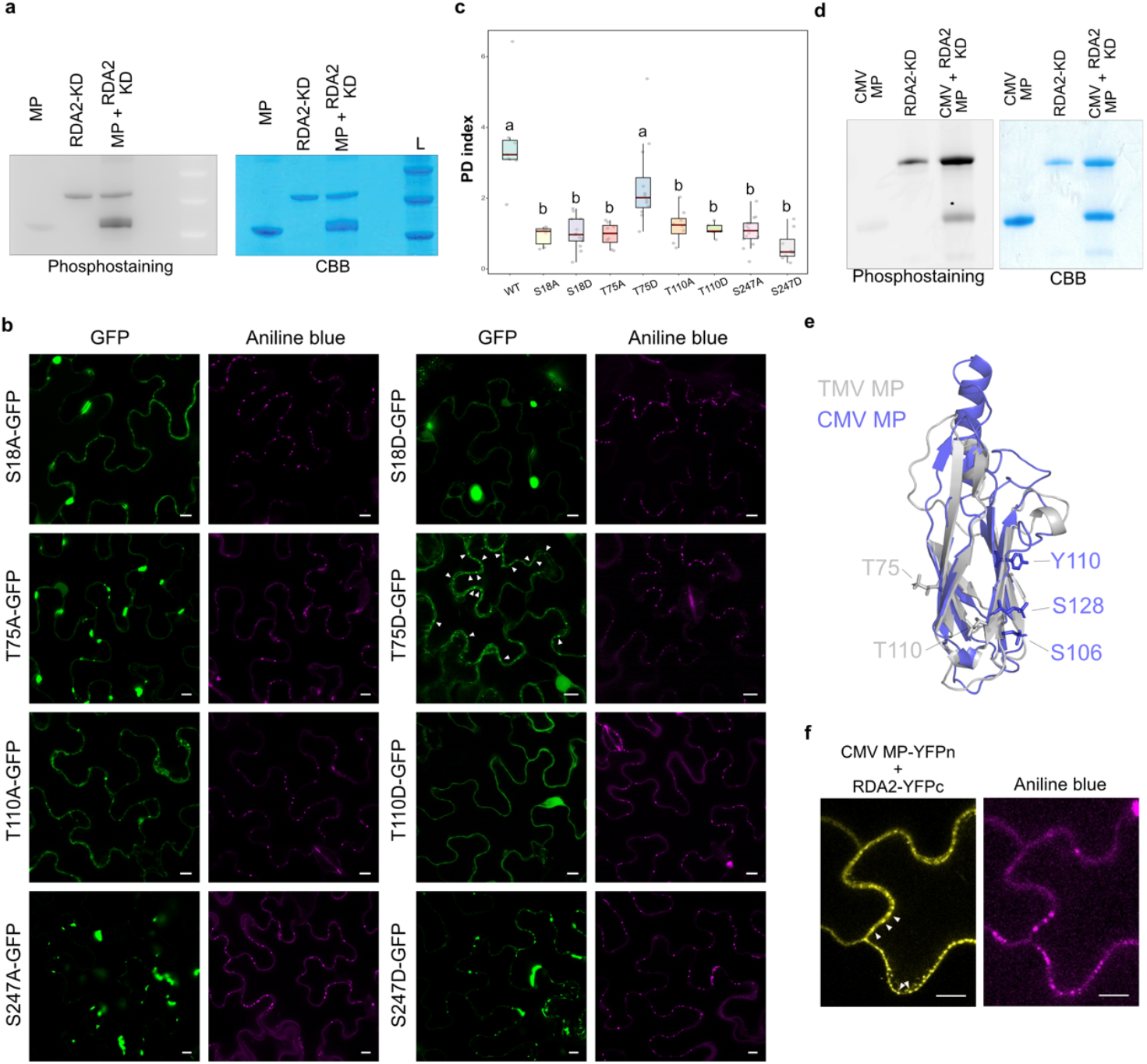
RDA2 phosphorylates 30K superfamily movement proteins from TMV and CMV, modulating plasmodesmatal association. **a** *In vitro* kinase assay showing phosphorylation of TMV MP by RDA2-KD. Phosphoprotein staining (left) indicates MP phosphorylation in the presence of RDA2-KD, and Coomassie brilliant blue (CBB) staining (right) confirms equal protein loading. **b** Subcellular localization of GFP-tagged TMV MP non-phosphorylatable mutants transiently expressed in *N. benthamiana* epidermal cells. All mutant variants displayed reduced PD targeting, showing cytoplasmic and nuclear accumulation, except for the phospho-mimetic T75D mutant, which retained punctate localization at plasmodesmata (arrowheads). Scale bars: 10 μm. **c** Quantitative analysis of plasmodesmal localization (PD index) showed that MP-T75D behaved similarly to the wild type, whereas the remaining mutants display significantly lower PD association. Different letters indicate significant differences according to Kruskal-Wallis test (p < 0.05) followed by Bonferroni post hoc comparison. At least 8 images from three biological replicates were analyzed per construct. **d** *In vitro* kinase assay showing phosphorylation of CMV MP by RDA2-KD, indicating that RDA2-mediated phosphorylation extends to movement proteins from different members of the 30K superfamily. **e** Structural superposition of predicted models of TMV MP (grey) and CMV MP (blue) showing conservation of the jelly-roll core characteristic of 30K superfamily movement proteins. Experimentally identified phosphorylation sites in TMV MP (T75, T110) and CMV MP (S106, Y110, S128) are mapped onto the structures, highlighting surface-exposed regions within the conserved fold. **f** Split-FP assay confirming interaction between RDA2 and CMV MP at PD in *N. benthamiana* epidermal cells. Arrowheads mark punctate fluorescence at plasmodesmata. Scale bars: 10 μm.

Finally, to test whether RDA2 directly interacts with MP, we performed *in vitro* pull-down assays using His-tagged recombinant proteins purified from *E. coli*. MP and RDA2-KD were incubated together and complexes were immunoprecipitated using anti-MP magnetic beads. Western blot analysis with anti-His antibodies and Coomassie blue staining revealed the presence of RDA2-KD in the eluate only when MP was included, confirming a direct interaction (**Fig. 2h)**. No RDA2-KD was detected in the absence of MP, demonstrating a direct and specific binding.

Collectively, these results demonstrate that RDA2 can interact with TMV MP both *in vitro* and *in planta*, and that the interaction is mediated by the kinase domain. Given that the cytoplasmic kinase domain of RDA2 mediates the interaction with MP, we hypothesized that RDA2 might directly phosphorylate the viral protein to modulate its function.

### RDA2 phosphorylates viral movement proteins and modulates their subcellular localization

RDA2 has been previously reported to activate mitogen-activated protein kinase (MAPK) 3 and 6 ^36^. To test whether RDA2 can also phosphorylate TMV MP, we performed *in vitro* kinase assays using purified recombinant proteins. TMV MP was incubated with RDA2-KD in the presence of ATP, and phosphorylation was detected by phosphoprotein gel staining. MP phosphorylation was observed only in the presence of RDA2-KD, whereas no signal was detected in control reactions lacking the kinase (**Fig. 3a**), demonstrating that RDA2 directly phosphorylates MP *in vitro*.

To identify specific phosphorylation sites, samples from the kinase reactions were analyzed by mass spectrometry. Four residues—S18, T75, T110, and S247—were consistently found to be phosphorylated (**Supplementary Table 2)**. To study their functional relevance, we generated phospho-dead (S/T>A) and phospho-mimetic (S/T>D) MP mutants by site-directed mutagenesis. GFP-tagged versions of wild-type and mutant TMV MPs were transiently expressed in *N. benthamiana* leaves and analyzed by confocal microscopy (**Fig. 3b**). Wild-type MP localized to PD, whereas phospho-dead mutants (S18A, T75A, T110A, S247A) failed to accumulate at PD and instead formed cytoplasmic aggregates and nuclear localization (**Fig. 3b)**. Most phospho-mimetic variants (S18D, T110D, S247D) also failed to restore PD targeting, indicating that mimicking a constitutive negative charge is insufficient to restore PD targeting at these positions (**Fig. 3b)**. Among all mutants, only MP-T75D retained PD localization comparable to the wild-type protein (**Fig. 3b**), indicating that a negative charge at T75 partially mimics the phosphorylated state required for PD association. This finding identified T75 as a key regulatory residue contributing to MP targeting and trafficking. Quantification of the PD index confirmed that MP-T75D is significantly enriched at PD (**Fig. 3c**). Expression of all constructs was verified by immunoblot analysis (**Supplementary Fig. 4)**.

**Figure 4.**
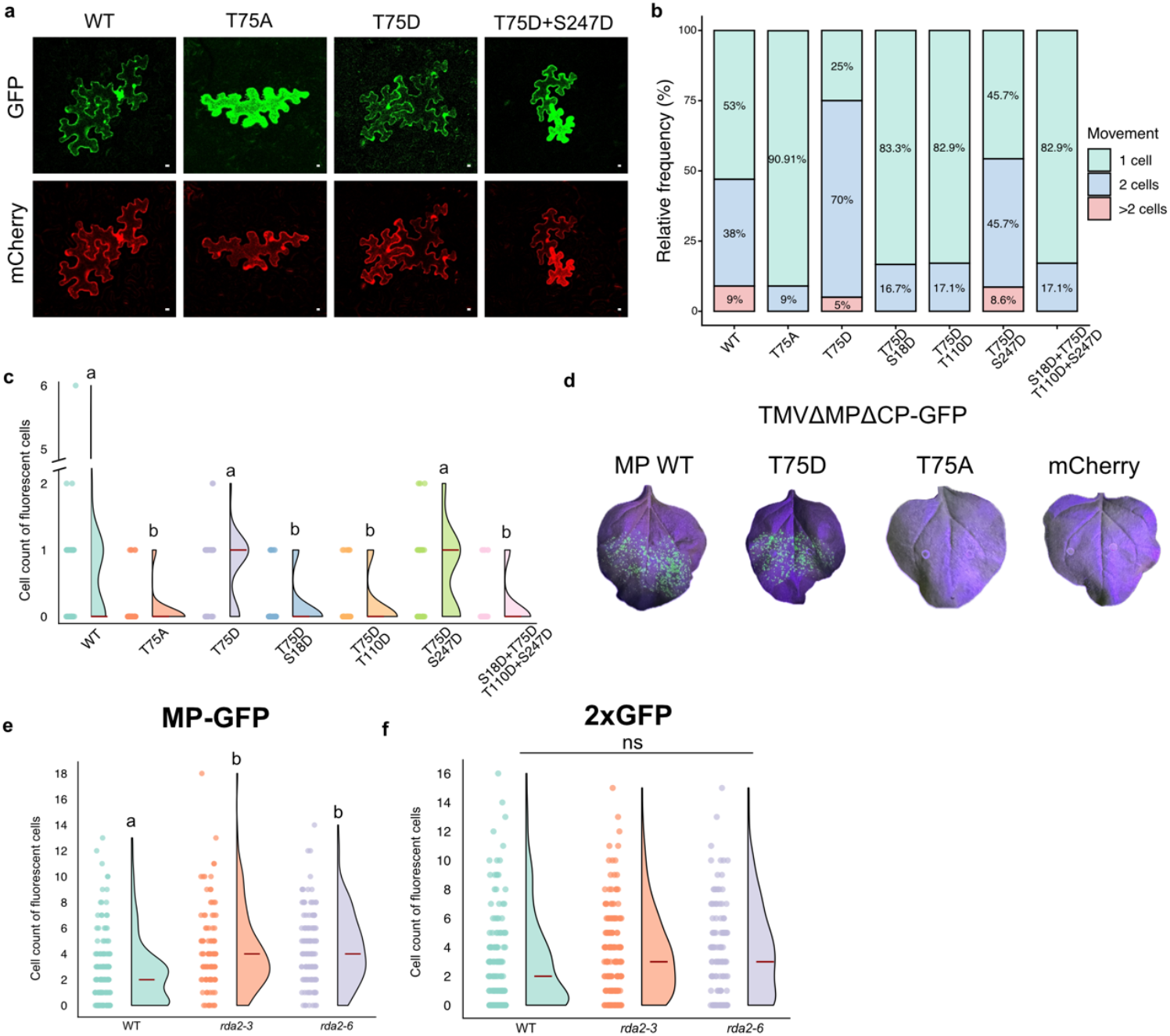
Phosphorylation at T75 determines cell-to-cell movement of TMV MP and is regulated by RDA2 *in planta*. **a** Intercellular transport assay in *N. benthamiana* leaves transiently expressing bicistronic constructs encoding MP-GFP variants together with a 3×mCherry tandem, which is immobile when expressed alone. MP-WT and MP-T75D promoted cell-to-cell spread of both reporters, whereas MP-T75A restricted fluorescence to the initially transformed cell. Among combinatorial phospho-mimetic mutants (S18D+T75D, T75D+T110D, T75D+S247D and the quadruple mutant), only T75D+S247D supported intercellular movement comparable to MP-WT. Scale bars: 50 μm. **b-c** Quantification of MP-mediated movement. **b** Frequency of multicellular fluorescence clusters for each MP variant. **c** Extent of spread measured as the number of fluorescent cells per cluster. Violin plots show median and interquartile range. Different letters indicate statistically significant differences (bootstrap analysis of the mean, p < 0.005). **d** Trans-complementation assay using TMVΔMPΔCP-GFP. The movement-deficient replicon is rescued by co-expression of MP-WT or MP-T75D, but not MP-T75A, as shown by GFP fluorescence in adjacent cells. Scale bar, 100 μm. **e** Microparticle bombardment of Arabidopsis leaves expressing MP-GFP. MP showed enhanced cell-to-cell spread in *rda2* mutant backgrounds compared with Col-0. Different letters indicate statistically significant differences (bootstrap analysis of the median, p < 0.005). Scale bar, 50 μm. **f** Control bombardment with 2×GFP (54 kDa) showed similar spread in Col-0 and *rda2*, indicating that differences observed in panel e were specific to MP mobility rather than global changes in PD permeability. Scale bar: 50 μm.

Together, these findings indicated that RDA2-dependent phosphorylation modulates TMV MP subcellular localization and highlight T75 as a major determinant of plasmodesmatal association.

### RDA2 interacts with and phosphorylates another member of 30K MP family

We next examined whether RDA2 activity is restricted to TMV MP or extends to additional members of the 30K movement protein family. Using CMV MP as a representative example, we performed *in vitro* kinase assays under identical conditions. RDA2 efficiently phosphorylated CMV MP, whereas no phosphorylation was detected in control reactions lacking the kinase (**Fig. 3d**), indicating that CMV MP is also a substrate of RDA2.

Mass spectrometry analysis identified multiple phosphorylation sites in CMV MP, including S73, Y75, S106, Y110 and S128 (**Supplementary Table 3**). Structural alignment of TMV and CMV MPs revealed conservation of the jelly-roll core ^16^ despite limited sequence similarity, while phosphorylated residues mapped to partially overlapping but non-identical surface-exposed regions (**Fig. 3e**). To test whether RDA2 also physically associates with CMV MP *in planta*, we performed split-fluorescent protein assays. Co-expression of RDA2 and CMV MP resulted in robust fluorescence reconstitution that partially overlapped with plasmodesmal markers (**Fig. 3f)**, indicating that RDA2 can interact with CMV MP at PD-associated sites.

Together, these results demonstrated that RDA2 can recognize and phosphorylate multiple 30K movement proteins, indicating that RDA2-mediated regulation of MP functionality may represent a broader antiviral mechanism rather than a TMV-specific effect.

### Phosphorylation at T75 is required for regulated MP cell-to-cell transport and PD gating

To understand the functional consequences of this phosphorylation on MP mobility, we next examined how the phospho-dead (T75A) and phospho-mimetic (T75D) MP variants behaved in intercellular transport assays.

We generated bicistronic constructs co-expressing either phospho-dead (T75A) or phospho-mimetic (T75D) mutants of MP and a tandem of three mCherrys (3xmCherry), which is immobile when expressed alone (**Supplementary Fig. 5)**, and transiently expressed in *N. benthamiana* epidermal cells. As expected, MP-WT-GFP and 3xmCherry moved between adjacent cells, reflecting the MP’s ability to gate PD (**Fig. 4a-c**). In contrast, MP-T75A-GFP and 3xmCherry were largely retained in the expressing cell, indicating that phosphorylation at T75 licenses MP for transport competence (**Fig. 4a-c**). Interestingly, the phospho-mimetic MP-T75D-GFP, which localizes to PD, supported intercellular movement and promoted 3xmCherry transport, while also accumulating in the cytosol and nucleus, in contrast to WT. This implicated that the presence of a negative charge at T75 is permissive for MP trafficking and PD localization but does not fully recapitulate the WT regulatory state (**Fig. 4a**). Because T75D was the only single phospho-mimetic variant that retained PD localization (**Fig. 3b,c**), we next examined whether additional phospho-mimetic substitutions modulate T75-dependent trafficking by analyzing combinatorial mutants. If MP trafficking were governed by an additive effect on phosphorylation, additional phospho-mimetic substitutions would be expected to maintain or enhance transport. We therefore analyzed a series of double (S18D+T75D, T75D+T110D, T75D+S247D) and quadruple phospho-mimetic mutants (S18D+T75D+T110D+S247D). Among these combinatorial variants, only T75D+S247D double mutant preserved intercellular movement comparable to WT and supported 3xmCherry transport, whereas all other combinations showed marked defects in movement (**Fig. 4a-c**). Quantification of cluster frequency and the number of cells traversed confirmed significant differences between WT, T75A, T75D and revealed that T75D+S247D is the only combinatorial mutant that restores WT-like trafficking (**Fig. 4b-c**). Consistent with this, independent localization analyses of the combinatorial phospho-mimetic mutants showed that only the T75D+S247D double mutant retained PD localization, whereas all other combinatorial mutants failed to accumulate at PD (**Supplementary Fig. 5**).

To determine whether the requirement for T75 phosphorylation observed in intercellular transport also applies during the infection, we used an MP-deficient TMVΔMPΔCP-GFP replicon in a trans-complementation assay with MP-WT, MP-T75A, or MP-T75D. Fluorescent foci were detected only when the replicon was complemented with MP-WT or MP-T75D, whereas MP-T75A failed to rescue movement, indicating that T75 phosphorylation was required for MP-mediated viral spread competence (**Fig. 4d**).

Because phosphorylation at T75 confers transport competence, we then asked whether the loss of RDA2 affects MP intercellular mobility *in planta*. MP-GFP (57 kDa) transport was analyzed after microparticle bombardment in Arabidopsis wild-type and two *rda2* mutants with independent alleles (*rda2-3* and *rda2-6*). MP-GFP spread more extensively in *rda2* mutants than in WT (**Fig. 4e**), indicating that RDA2 normally constrains MP intercellular movement *in planta*. As a control, tandem GFP fusion (2×GFP; 54 kDa) showed no significant differences between genotypes, confirming that the observed effect is specific to MP and not due to a general change in PD permeability (**Fig. 4f**).

Together, these results identify T75 phosphorylation as a key regulatory checkpoint for MP-mediated plasmodesmatal transport and viral cell-to-cell movement. Loss of RDA2 enhanced MP intercellular mobility in planta, supporting a role for RDA2-mediated phosphorylation in constraining MP trafficking.

### RDA2 act as antiviral host factor restricting TMV infection

Having established that RDA2 directly phosphorylates TMV MP and restricts its plasmodesmal mobility, we next asked whether this regulation impacts viral spread. To evaluate the physiological relevance of RDA2 during TMV infection, we quantified viral accumulation in Arabidopsis Col-0 and the two independent *rda2* mutant alleles (*rda2-3* and *rda2-6*).

Plants were mechanically inoculated with TMV, and systemic, non-inoculated leaves were harvested at 7 days post inoculation (dpi) for RT-qPCR analysis of viral RNA. TMV accumulation in Col-0 plants showed strong variability, with several individuals displaying undetectable or very low viral RNA levels, consistent with the low susceptibility of Arabidopsis to TMV ^37^. In contrast, both *rda2-3* and *rda2-6* mutants exhibited a higher frequency of infected plants and increased viral RNA accumulation (Fig. 5a). When infection outcomes were categorized based on viral RNA levels, *rda2* mutants displayed a clear shift toward the high-accumulation class compared to Col-0 (Fig. 5b), indicating enhanced systemic infection in the absence of RDA2.

We next analyzed TMV coat protein (CP) accumulation by immunoblotting in both inoculated and systemic leaves. CP levels in inoculated leaves were comparable between Col-0 and *rda2-3* plants, indicating that early stages of infection and local viral accumulation were not strongly affected by loss of RDA2 (Fig. 5c). In contrast, CP accumulation in systemic leaves was markedly reduced in Col-0 plants but readily detectable in *rda2-3* mutants (Fig. 5c), showing that loss of RDA2 preferentially enhances systemic viral accumulation rather than local infection.

These infection phenotypes are consistent with a role for RDA2 in restricting viral spread. While TMV accumulation in inoculated leaves was comparable between genotypes, *rda2* mutants showed enhanced viral accumulation in systemic tissues (Fig. 5a-c), indicating that RDA2 preferentially limits systemic infection.

**Figure 5.**
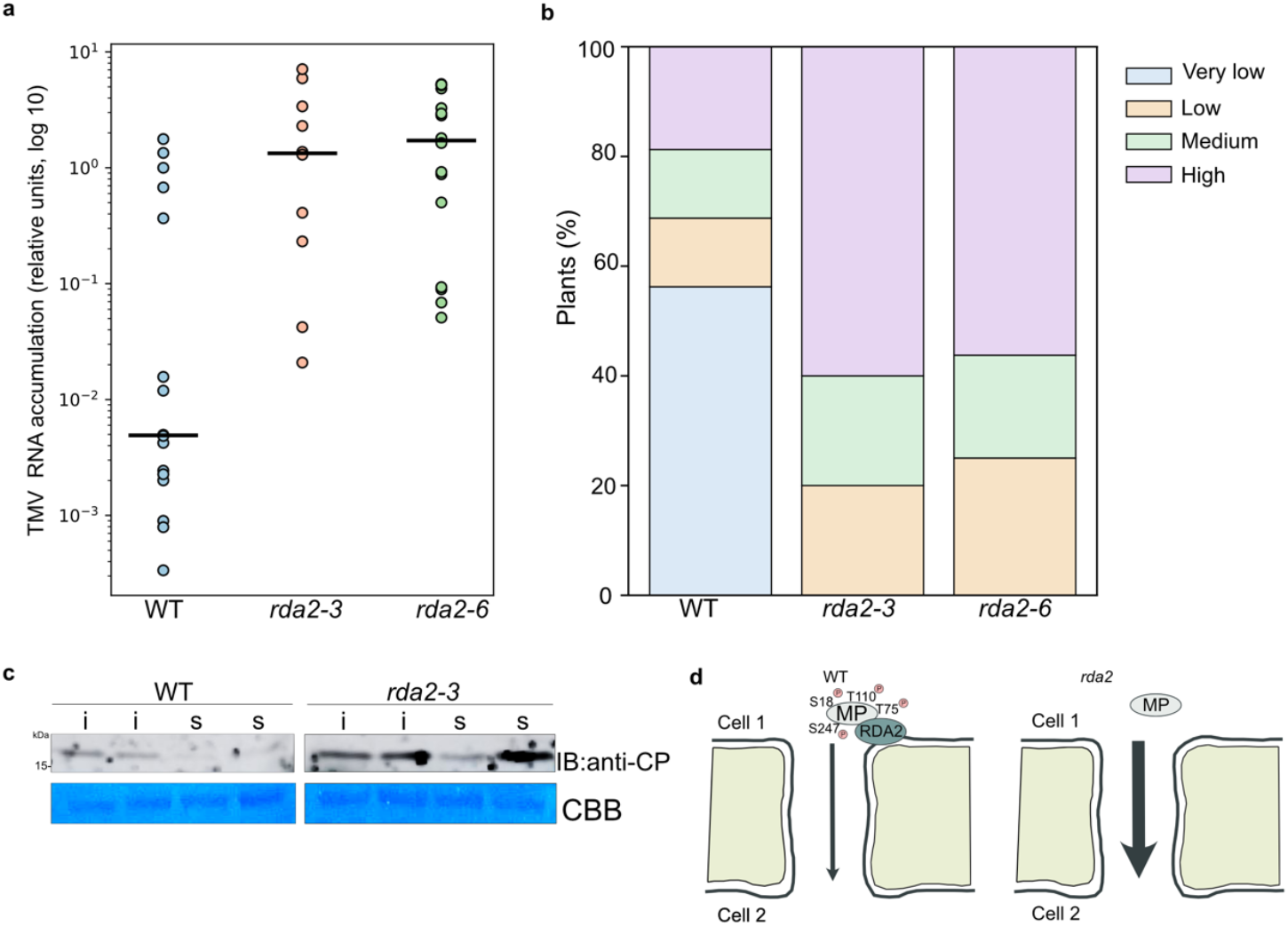
RDA2 restricts systemic TMV infection by limiting viral movement. **a** TMV RNA accumulation in systemic leaves of Arabidopsis *rda2* mutants. TMV RNA levels were quantified by RT-qPCR in systemic, non-inoculated leaves of Arabidopsis Col-0, *rda2-3* and *rda2-6* plants at 7 dpi. Each dot represents an individual plant, and horizontal bars indicate the median. Data are shown on a logarithmic scale. Statistical analysis using a Kruskal-Wallis test revealed a significant difference among genotypes (p < 0.001). Posthoc pairwise comparisons using the Mann-Whitney U test showed significantly higher TMV RNA accumulation in both *rda2-3* and *rda2-6* compared to Col-0 (p < 0.01). **b** Distribution of systemic infection outcomes in Col-0 and *rda2* mutants. The RT-qPCR data shown in **a** were grouped into infection severity classes and plotted as the percentage of plants per category for each genotype. Severity classes were defined as follows: very low (< 0.01), low (0.01-0.1), medium (0.1-1) and high (> 1) TMV RNA accumulation, expressed as relative quantities (RQ). *rda2* mutants display a marked shift toward higher accumulation classes compared to Col-0, indicating enhanced systemic infection in the absence of RDA2. **c** Immunoblot analysis of TMV CP accumulation in inoculated (i) and systemic (s) leaves of Arabidopsis Col-0 and *rda2-3* plants at 7 dpi. Total protein extracts were probed with anti-CP antibodies. Coomassie Brilliant Blue (CBB) staining of the membrane is shown as a loading control. **d** In wild type plants, RDA2 localizes to plasmodesmata and phosphorylates the TMV MP. Phosphorylation at T75 is required for MP plasmodesmal targeting, whereas additional phosphorylation at S18, T110 and S247 limits MP intercellular transport, thereby restricting cell-to-cell and long-distance viral spread. In the absence of RDA2, loss of MP phosphorylation enhances plasmodesmal transport and promotes systemic infection.

## Discussion

During the process of pathogen coevolution, plants have evolved sophisticated regulatory layers to control immune activation while preserving essential intercellular communication. While PD are crucial for developmental signaling, they also constitute a vulnerability that viruses exploit for cell-to-cell movement. Despite extensive work on viral MPs and host factors that modulate PD structure, the contribution of plasma-membrane receptor-like kinases (RLKs) to antiviral PD regulation has remained largely unexplored. Here, we show that the lectin RLK RDA2 directly phosphorylates viral MPs at plasmodesmata, revealing an unanticipated receptor-level mechanism that modulates intercellular connectivity during infection.

Using proximity labeling, we found consistently RDA2 in the immediate vicinity of the TMV MP *in vivo*, together with known MP interactors such as SYTA ^32^. Proximity labeling has recently emerged as a powerful approach to capture transient plant-virus interactions ^38,39^, and our findings extend its application by identifying a plasma membrane receptor-like kinase as a stable component of the MP-associated proteome. The recovery of RDA2 together with established MP-associated proteins supports the emerging view that plasmodesmata are organized into discrete plasma membrane nanodomains enriched in specific regulatory factors. Previous work has shown that PD-associated proteins segregate into specialized membrane subdomains that function as signaling platforms rather than uniform conduits ^40^. Our findings extend this concept by positioning a plasma membrane receptor-like kinase within MP-associated PD nanodomains, suggesting that viral movement is locally regulated by receptor-mediated signaling at PD.

Beyond RDA2, our proximity labeling dataset recovered two additional signaling components with potential relevance to MP regulation at PD: a glycine-rich protein (AT4G02450) and the calcium-responsive kinase CIPK24/SOS2 (AT5G35410). The presence of a Ca^2+^-regulated kinase is notable given the central role of Ca^2+^ fluxes in PD gating, as demonstrated in multiple systems where cytosolic Ca^2+^ elevations rapidly reduce PD permeability ^41–44^. Although the functional contribution of these candidates remains to be determined, their identification suggests that MP-associated nanodomains may integrate receptor- and Ca^2+^-dependent signaling pathways to dynamically control PD function during infection.

The identification of kinases within the MP-associated proteome suggests that phosphorylation is an integral component of MP regulation at plasmodesmata. RDA2-mediated modification appears to control a defined regulatory checkpoint required for PD targeting and intercellular transport. Notably, one of the modified residues (S247) lies within a region previously implicated in phosphorylation-dependent movement control ^45^. These findings refine longstanding models in which MP phosphorylation acts as a molecular switch controlling transitions between RNA-binding, trafficking and release-competent states ^46,47^, by showing that a plasma-membrane RLK directly imposes one of these transitions at PD.

Together, these findings indicate that MP transport competence is not governed by a simple on/off phosphorylation switch, but instead depends on a coordinated, multi-site phosphorylation state. Phosphorylation at T75 acts as a key licensing event for PD targeting, while additional RDA2-dependent modifications modulate MP mobility, as supported by the behavior of combinatorial mutants. This type of balance-based regulation is increasingly recognized in plant signaling pathways, suggesting that similar principles may operate to control viral movement at plasmodesmata ^48^. Our findings extend this regulatory principle to virus-host interactions, demonstrating that balanced phosphorylation of a viral movement protein can similarly determine transport competence at PD.

Importantly, this phosphorylation-dependent regulation has clear consequences at the level of viral infection in planta. Our functional assays establish RDA2 as an antiviral host factor that restricts viral spread in planta by limiting MP-mediated intercellular movement. MP-GFP spreads more extensively in rda2 mutants than in wild-type plants, and TMV accumulates to higher levels in rda2 backgrounds, indicating that RDA2 dampens systemic infection. Among plasma membrane RLKs, NIK1 represents the best-characterized antiviral example, acting through a translation-suppression pathway that restricts viral accumulation ^49,50^. Although mechanistically distinct, the comparison with NIK1 highlights how different RLKs can restrict viral infection at distinct regulatory layers, from translation control to cell-to-cell movement. Together, these observations support a model in which RDA2 functions at plasmodesmata to restrict systemic viral spread by phosphorylating the viral movement protein and limiting its intercellular mobility. In this framework, RDA2-mediated phosphorylation reduces the efficiency with which local infection foci give rise to long-distance viral spread, whereas loss of RDA2 disrupts this regulatory checkpoint, resulting in enhanced MP trafficking through plasmodesmata and increased systemic infection (Fig. 5d).

The ability of RDA2 to interact with and phosphorylate the CMV MP may indicate that this regulatory mechanism extends beyond TMV and is likely conserved across members of the 30K MP superfamily. While most phosphorylation sites diverge between TMV and CMV MPs, structural alignment revealed that the region surrounding TMV T110 spatially coincides with a phosphorylated residue in CMV MP. This observation raises the possibility that phosphorylation targets a conserved regulatory surface within the jelly-roll core, even though the precise phospho-acceptor residues have diverged during evolution.

Consistent with a broader and potentially conserved role for RDA2-like kinases in antiviral defense, single-cell transcriptomic analyses in *Nicotiana tabacum* revealed that multiple RDA2 orthologs (Nta03g29160, Nta04g21900 and Nta17g14590) undergo infection-dependent transcriptional changes during TMV and CMV infection ^51^. These orthologs display distinct induction or repression patterns across cell types, indicating that RDA2-like kinases participate in antiviral transcriptional programs in a natural tobamovirus host. This cross-species evidence reinforces the idea that phosphorylation-based regulation of 30K MPs may represent a conserved antiviral module across angiosperms.

Together, our findings fit into a broader emerging paradigm in which plasmodesmata are viewed as immunologically active microdomains whose permeability is dynamically regulated through coordinated activities of plasma-membrane receptors, membrane scaffolds, and callose-modifying enzymes ^52–54^. Such mechanisms collectively shape the PD environment and influence viral movement. Our work positions RDA2 within this regulatory landscape, identifying it as a receptor-like kinase that directly modifies a viral movement protein at PD. By positioning a receptor-like kinase upstream of MP phosphorylation, our work establishes a mechanistic link between immune receptor signaling and the local control of PD transport during viral infection.

In summary, our study reveals that the receptor-like kinase RDA2 restricts viral spread by directly phosphorylating 30K MPs at PD and modulating their trafficking behavior. By linking immune receptor function with local post-translational control of PD transport, RDA2 defines a previously uncharacterized antiviral mechanism and offers a conceptual framework for engineering broad-spectrum resistance through manipulation of RLK-MP regulatory modules.

## Methods

### Plant material

*Arabidopsis thaliana rda2-6* (G585R) mutant and the T-DNA insertion mutant *rda2-3* (SALK_143489C) were kindly provided by Dr. Kato (University of Kyoto) ^29^. UBQ10:MP-TbID-mVenus mutant line was generated by floral dipping of Col-0 ecotype plants. For microparticle bombardment, Arabidopsis Col-0, *rda2-3*, and *rda2-6* seedlings were grown on half-strength Murashige and Skoog (MS) medium supplemented with 1% (w/v) sucrose and solidified with 0.8% agar plates for 4 weeks in short day conditions (10 h light, 14 h darkness) at 23 °C with a light intensity to 130 μmol m-2 s–1. For susceptibility assays, Arabidopsis plants were grown on soil in an environment-controlled chamber at 16 h light/8 h darkness, 23 °C, ∼130 *μ*mol m−2 s−1, 60% humidity for 3 weeks. N*icotiana benthamiana* plants were grown on soil in a growth-controlled chamber (16 h light/8 h darkness, 23 °C, ∼90 *μ*mol m−2 s−1, 60% humidity) for 3 weeks.

### Plasmid construction

To generate UBQ10:MP-TbID-mVenus, TMV MP (GenBank accession AJ509081) was synthesized with *BsaI* sites and cloned into plasmid pGGC000 to generate module C. UBIQUITIN10 promoter and TbID-mVenus were used in module A and D, respectively. All modules were transferred to binary vector pGGZ003 by GreenGate reaction ^55^. For subcellular localization and split-FP, TMV MP, CMV MP (strain fny, GenBank D10538) and RDA2 (AT1G11330) ORFs were PCR-amplified with *BsaI* sites and ligated into *BsaI*-digested pGGC000 plasmid. Non-phosphorylatable and phosphorylatable TMV MP mutants were generated by site-directed mutagenesis. A ß-estradiol-inducible promoter was used in module A. GFP, mCherry, nYFP or cYFP were used as module D. For cell-to-cell transport assays, bicistronic plasmids were generated with the corresponding TMV MP-GFP fusion version either WT, T75A or T75D cloned into pGGM and 3xmCherry tandem cloned into pGGN under 35S and UBQ10 promoters, respectively. Subsequently, pGGM-35S-MP-WT, pGGM-35S-MP-T75A or pGGM-35S-MP-T75D; and pGGN-UBQ10-3xmCherry were subcloned into binary vector pGGZ003 by restriction cloning with *BsaI*. The TMVΔMPΔCP-GFP replicon was generated as described in Borniego et al., 2016. Briefly, the MP ORF was truncated at positions 57-545 by PCR in pJL-TRBO-G vector ^56^. For recombinant protein expression, TMV MP, CMV MP and RDA2 KD ORFs were cloned into a pET30a (Novagen) expression vector, to generate N-terminal 6xHis-tagged proteins. All constructs were verified by Sanger sequencing.

### Agroinfiltration

*Agrobacterium tumefaciens* cultures carrying expression constructs were grown to appropriate OD600, pelleted, resuspended in infiltration buffer (10 mM MES pH 5.6, 10 mM MgCl2, 150 μM acetosyringone) and co-infiltrated into *N. benthamiana* leaves. For split-FP and localization assays, typical final OD600 values were 0.3 for expression constructs and 0.1 for P19 silencing suppressor unless otherwise stated. Samples were imaged 2-4 days post-infiltration.

### Sample preparation for LC-MS/MS

Arabidopsis plants were cultivated in a hydroponic culture system as described in Schlesier et al. (2003) ^57^ for root collection. Approximately 25 mg of surface-sterilized seeds were placed on mesh within glass jars containing 150 mL of ½ MS (Murashige and Skoog, Sigma-Aldrich) supplemented with 0.5% sucrose, and grown on a shaker in a growth chamber (16/8 h day/night, 22 °C, 110 ± 10 μE s^−1^ m^−2^) for 22 days. The growth medium was then replaced with 50 μM biotin (Sigma-Aldrich) and plants were incubated for 1 h with gentle shaking at room temperature. Plants were washed with ice-cold water and tissue was frozen in liquid nitrogen. Protein extraction was performed according to Mair et al. (2019). Approximately 1.5 g of tissue was ground to a fine powder and resuspended in ice-cold extraction buffer (50 mM Tris pH 7.5, 150 mM NaCl, 0.1% SDS, 1% Triton X-100, 1 mM EGTA, 1 mM DTT, 0.5% sodium deoxycholate, 1 mM PMSF, and 1× complete protease inhibitor) and incubated on a rotor wheel at 4 °C for 10 min.

Lysozyme (1 μL, 1 mg/mL) was added and samples were incubated for an additional 15 min at 4 °C. Samples were sonicated four times for 30 s in an ice bath with 1.5 min intervals on ice. Lysates were centrifuged at 15,000 × g and the supernatant was subjected to biotin removal using Cytiva PD-10 desalting columns following the manufacturer’s instructions. Protein concentration was determined by Bradford assay (Bradford, 1976). For streptavidin enrichment, 2 mg of total protein per sample was incubated with 80 μL streptavidin magnetic bead slurry (Streptavidin Mag Sepharose, Cytiva 28-9857-38; beads pre-washed with 500 μL extraction buffer) supplemented with PMSF and protease inhibitor. Samples were incubated overnight at 4 °C on a rotor wheel. Beads were collected using a magnetic rack and washed once each with the following solutions: cold extraction buffer, cold 1 M KCl, cold 100 mM Na_2_CO_3_, 2 M urea in 10 mM Tris pH 8 at room temperature, and finally cold extraction buffer. Beads were spun down and the remaining washing buffer was removed.

### On-bead trypsin digestion

Beads were washed with 1 ml 50 mM Tris, pH 7.5, transferred to a new tube, washed again with 1 ml 50 mM Tris, pH 7.5, and then washed twice with 2 M urea in 50 mM Tris, pH 7.5. The washing buffer was then replaced with 80 μl trypsin buffer (50 mM Tris pH 7.5, 1 M urea, 1 mM DTT, 0.4 μg trypsin). Samples were incubated at 37°C for 3 h. The supernatant was collected in a new tube, and the beads were washed twice with 60 μl 1 M urea in 50 mM Tris pH 7.5, all supernatants were combined. DTT was added to a final concentration of 4 mM and samples were incubated at 25°C for 30 min with shaking. Iodoacetamide was then added to a final concentration of 10 mM and samples were incubated at 25°C for 45 min with shaking. An additional 0.5 μg trypsin was added for overnight digestion (14.5 h) at 37°C. The reaction was stopped by acidification with TFA to pH ≤ 2.

Prior to LC-MS/MS analysis, the peptides were desalted over C18-Sage tips as described ^58,59^.

### LC-MS/MS analysis of peptides

Peptide mixtures were analyzed by nanoflow Easy-nLC (Thermo Scientific) and Orbitrap hybrid mass spectrometer (Q-Exactive HF, Thermo Scientific). Peptides were eluted from a 75 μm x 25 cm analytical C18 column (PepMan, Thermo Scientific) on a linear gradient running from 5% to 90% acetonitrile for 70 min. Proteins were identified based on the information-dependent acquisition of fragmentation spectra of multiple charged peptides. Up to ten data-dependent MS/MS spectra were acquired for each full-scan spectrum acquired at 60,000 full-width half-maximum (FWHM) resolution.

### Peptide and protein identification

Protein identification and ion intensity quantitation was carried out by MaxQuant version 2.0.3.0 ^60^. Spectra were matched against the Arabidopsis proteome (Araport11_genes.201606.pep.fasta, 48359 entries) using Andromeda ^61^. Thereby, carbamidomethylation of cysteine was set as a fixed modification; oxidation of methionine as well as phosphorylation of serine, threonine and tyrosine were set as variable modifications. Mass tolerance for the database search was set to 20 ppm on full scans and 0.5 Da for fragment ions. Multiplicity was set to 1. For label-free quantitation, retention time matching between runs was chosen within a time window of two minutes. Peptide false discovery rate (FDR) and protein FDR were set to 0.01, while site FDR was set to 0.05. Hits to contaminants (e.g. keratins) and reverse hits identified by MaxQuant were excluded from further analysis.

### Label-free peptide and protein quantification

We followed a label-free quantitative approach based on LFQ values as quantitative information obtained from MaxQuant ^62^. For protein identification and quantitation, protein groups information (protein groups.txt) was used. Data analysis was performed using Perseus software and log2-transformation of the raw LFQ-values was used ^63,64^.

### Statistical analysis and data visualization for protein identification

MAPMAN was used for protein functional classification ^65^. SUBA4 was used for subcellular location identification ^66^. Detailed protein function was manually updated with the support of TAIR ^67^. Other statistical analyses were carried out with ORIGIN PRO software (version 2022b) and Excel (Microsoft, 2019). The corresponding data are presented in the supplemental data.

### Confocal microscopy

Confocal microscopy was performed on a Leica Stellaris 8 microscope (Leica Microsystems) using an HC PL APO CS2 63x/1.30 glycerol immersion objective. To visualize pit fields, plant leaves were infiltrated with 0.01% (w/v) aniline blue (Sigma Aldrich) solution (PBS buffer, pH 8.0). Images were captured sequentially with excitation at 405 nm (aniline blue), 480 nm (GFP), 514 nm (mVenus) or 561 nm (mCherry) and emission was collected at 440-480 nm (aniline blue), 500-550 nm (GFP), 520-570 nm (mVenus) or 590-610 nm (mCherry). Bombardment analysis was performed on a Zeiss LSM 900 (Carl Zeiss, Oberkochen, Germany) equipped with Airyscan GaAsP-PMT detectors and diode lasers using a 40x/1.20 water immersion objective (CApochromat 40x/1.20 W Korr FCS). GFP was excited at 488 nm and collected in a 500-550 nm window.

### Purification of recombinant proteins from *E. coli*

Arabidopsis RDA2 KD has a predicted nucleoid-binding function. To prevent nucleic acid contamination and misfolding, the protein was expressed in *E. coli* and purified in the presence of nuclease. Cells were harvested by centrifugation at 4,000 × g for 15 min at 4 °C and resuspended in lysis buffer (50 mM Tris-HCl, 500 mM NaCl, 1% Triton X-100, pH 8.0). Cells were lysed by sonication, and Benzonase® (250 U/μL; Merck) was added and incubated 30 min at 4 °C to degrade bacterial DNA/RNA. Lysates were clarified by centrifugation at 20,000 × g for 30 min at 4 °C. The soluble fraction was applied to Ni^2+^-NTA resin (GenScript) and washed with equilibration buffer (50 mM Tris-HCl, 500 mM NaCl, pH 8.0) containing up to 500 mM imidazole to reduce nucleic-acid contamination. RDA2 KD was eluted stepwise (20/50/500 mM imidazole in 50 mM Tris-HCl, 500 mM NaCl, pH 8.0), concentrated, and polished by size-exclusion chromatography in SEC buffer (50 mM Tris-HCl, 150 mM NaCl, 10% glycerol, pH 8.0). Removal of nucleic acids was confirmed by DNA detection assay (PicoGreen; Invitrogen). His-tagged TMV MP and CMV MP were purified from inclusion bodies as described in ^68,69^. Protein expression was induced in *E. coli* at 15 °C for 16 h with 0.5 mM IPTG. Cells were harvested by centrifugation at 4,000 × g for 15 min at 4 °C and resuspended in lysis buffer (50 mM Tris-HCl, 150 mM NaCl, 1% Triton X-100, pH 8.0), then lysed by sonication. Inclusion bodies were collected by centrifugation at 20,000 × g for 30 min at 4 °C, washed, and solubilized in inclusion body solubilization buffer (50 mM Tris-HCl, 7 M guanidine hydrochloride, pH 8.0). The solubilized protein was clarified by centrifugation (20,000 × g, 30 min, 4 °C) and applied to Ni^2+^-NTA resin (GenScript) equilibrated in equilibration buffer (50 mM Tris-HCl, 8 M urea, pH 8.0). Bound protein was eluted stepwise (20/50/500 mM imidazole in 50 mM Tris-HCl, 8 M urea, pH 8.0) and refolded by stepwise dialysis into refolding buffer (50 mM Tris-HCl, 500 mM NaCl, 10% glycerol, pH 8.0). Protein purity was confirmed by SDS-PAGE.

### *In vitro* phosphorylation assays

*In vitro* phosphorylation assay was performed as described ^22^ with minor modifications. Briefly, 1 μg of TMV MP and 0.4 μg of KD were assembled in a kinase reaction buffer containing 50 mM HEPES (pH 7.5), 10 mM MgCl_2_, 2 mM DTT, 2 mM EGTA, and 2.2 mM CaCl_2_ and kept on ice until ATP was added to a final concentration of 2 mM. The reaction mixture was immediately transferred to 30 °C and incubated for 20 min. The reaction was stopped by adding 1/4 volume of SDS-PAGE 4× loading buffer and boiling for 5 min. The 20-μl aliquots of the reaction mixtures were electrophoretically resolved on a 12% SDS-PAGE gel, followed by staining with the Pro-Q diamond phosphoprotein gel stain according to manufacturer instructions (Fisher Scientific).

The total protein in the phosphostained gel was detected by subsequent staining with Coomassie brilliant blue. Protein product images were recorded using an Amersham Imager 680 (GE Healthcare Life Sciences).

### Identification of phosphorylated residues

Samples were digested with the following standard procedure. Protein products (~2×2 mm) were excised from gels, destained, and washed with MilliQ water followed by ammonium bicarbonate buffer in 50% acetonitrile. After drying, proteins were reduced with DTT (20 min, 56 °C) and alkylated with iodoacetamide (30 min, RT, dark). Samples were then washed again with buffer and buffer/acetonitrile, dried, and incubated in trypsin solution (0.5 μg Trypsin Gold in ammonium bicarbonate buffer with 0.01% ProteaseMax; Promega) for 10 min at 4 °C. Prior to the trypsin digestion, samples were enriched in phosphorylated proteins using the Pierce ™ Phosphoprotein Enrichment Kit according to the manufacturer instructions. Digestion proceeded overnight (16 h, 37 °C). Peptides were recovered by sequential washes with acetonitrile/TFA and acetonitrile, with all supernatants pooled and dried under vacuum. Dried peptide samples were resuspended in 20 μl of buffer A (water/acetonitrile/formic acid, 94.9:5:0.1) and analyzed using an Agilent 1290 Infinity II HPLC system coupled to an Agilent 6550 Q-TOF mass spectrometer equipped with an AJS-Dual ESI interface, operated with MassHunter Workstation (Rev. B.08.00). Samples were loaded onto an Agilent AdvanceBio Peptide Mapping column (100 × 2.1 mm, 2.7 μm) thermostatted at 50 °C and run at a flow rate of 0.4 ml/min. After injection, the column was washed with buffer A for 3 min, followed by peptide elution using a linear gradient of 0-40% buffer B (water/acetonitrile/formic acid, 10:89.9:0.1) over 40 min, 40-95% B for 8 min, held at 95% B for 3 min, and re-equilibrated for 6 min. The mass spectrometer was operated in positive ion mode with the following parameters: nebulizer pressure, 35 psi; drying gas, 14 l/min at 300 °C; sheath gas, 11 l/min at 250 °C; capillary, 3500 V; nozzle, 100 V; fragmentor, 360 V; octopole RF Vpp, 750 V. Data were acquired in extended dynamic range mode (4 GHz) with MS scans from m/z 50-1700 at 8 spectra/s and MS/MS scans at 3 spectra/s. Auto MS/MS was enabled with abundance-based precursor selection (up to 20 precursors per cycle), dynamic exclusion after two spectra, and ramped collision energy (slope 3.68, offset -4.28). Spectra were processed in Spectrum Mill MS Proteomics Workbench (Rev. B.06.00.201, Agilent Technologies). Extraction parameters included unmodified or carbamidomethylated cysteines, precursor [M+H]+ range 50-10000 m/z, maximum charge +5, and a minimum MS signal-to-noise ratio of 25. Database searches were performed against the relevant protein database with trypsin specificity (up to 5 missed cleavages), fixed carbamidomethylation of cysteines, and variable modifications including STY phosphorylation, methionine oxidation, and N-terminal pyroglutamate formation. Search tolerances were set to 20 ppm for precursor ions and 50 ppm for product ions. Peptide and protein identifications were validated by automatic thresholds using reversed database scoring.

### Virus material and inoculation

Tobacco mosaic virus (TMV; PV-1252) was obtained from the DSMZ collection. For inoculum preparation, *N. benthamiana* leaves systemically infected with TMV were homogenized in inoculation buffer (0.1 M Na_2_HPO_4_, 0.5 M NaH_2_PO_4_, pH 7.4). Two-week-old Arabidopsis plants were mechanically inoculated by gently rubbing 20 μL of the clarified extract onto the fourth and fifth true leaves.

### RNA extraction and RT-qPCR

At 7 days post inoculation (dpi), upper, non-inoculated Arabidopsis leaves were individually harvested from each plant. Total RNA was extracted using NZYol® (nzytech, Lisbon, Portugal), followed by phenol-chloroform purification and ethanol precipitation. The purified RNAs were treated with DNase I (Sigma-Aldrich), quantified using a NanoDrop One spectrophotometer (Thermo Fisher Scientific), and assessed for integrity by agarose gel electrophoresis. Relative quantification of TMV RNA was performed by one-step reverse transcription quantitative PCR (RT-qPCR) using the NZYSpeedy One-step RT-qPCR Green ROX Plus kit (NZYTech), following the manufacturer’s instructions. TMV-specific primers (Supplementary Table S4) were used for viral RNA amplification. Arabidopsis ACTIN2 (AT3G18780) was used as an internal reference gene for normalization, and relative TMV accumulation was calculated using the ΔΔCt method.

### *In vitro* interaction assays

Co-immunoprecipitation (Co-IP) assays were performed using TMV-MP-6xHis and RDA2-KD-6xHis recombinant proteins. Briefly, 2 μg of anti-MP antibody (Alpha Diagnostic) were pre-incubated with 20 μL of Protein A magnetic beads (Thermo Fisher Scientific) in 200 μL of interaction buffer (Thermo Fisher Scientific) for 1 h at 4 °C with gentle rotation. In parallel, 1.5 μg of MP-6xHis were mixed and incubated with 2 μg of KD-6xHis for 1 h at 4 °C in kinase buffer with or without ATP. The protein mixture was then added to the antibody coated beads and incubated for an additional 1 h at 4 °C. Beads were washed three times with washing buffer, and bound proteins were eluted with elution buffer and boiled in 4x Laemmli sample buffer. Samples were analyzed by western blotting with anti-His (Thermo Fisher Scientific) and Coomassie brilliant blue staining. Negative controls included individual proteins incubated separately.

### PD index quantification

Plasmodesmal enrichment of fluorescently tagged proteins was quantified using the PD index as previously described (Grison et al., 2019). To minimize user-dependent bias, analyses were performed using our recently developed automated PD index analysis pipeline (Gombos et al., 2023). Briefly, clearly identifiable PD were selected, and fluorescence channels were separated for analysis. Plasmodesmata were detected by automated thresholding in Fiji using the YEN algorithm, with minor user adjustments when required. The centroid coordinates of individual plasmodesmata were subsequently determined, and square regions of interest (ROIs) were defined around each site. The PD index was calculated as the ratio between the mean fluorescence intensity within PD ROIs and the mean intensity of background ROIs. Background ROIs were defined in regions displaying low aniline blue signal, and areas previously identified as plasmodesmata were automatically excluded from background ROI placement using image-based processing.

### Statistical analysis

Statistical analyses were performed using GraphPad Prism 10 unless otherwise stated. PD index was calculated. For PD index and related localization quantifications, statistical significance among multiple groups was assessed using a Kruskal-Wallis test followed by Bonferroni-corrected post hoc comparisons. Data are presented as boxplots showing the median and interquartile range. For cell-to-cell movement assays in *N. benthamiana* and for microparticle bombardment assays in *Arabidopsis*, statistical differences between constructs/genotypes were evaluated using bootstrap-based inference implemented in RStudio, following the approach described by Matthew et al. Specifically, bootstrap analysis of the mean (movement assay quantifications) or of the median (bombardment assays) was applied as indicated in the figure legends, using a significance threshold of p < 0.005.

### Transport assays

#### MP cell-to-cell movement assays

*A. tumefaciens* cultures containing bicistronic constructs expressing either WT, T75A or T75D fused to GFP and 3xmCherry were agroinfiltrated into *N. benthamiana* leaves at OD_600_ 0.0001 combined with an *A. tumefaciens* strain containing the P19 silencing suppressor (OD_600_ 0.1). Samples were visualized 3 dpi under confocal microscope. The number of cells expressing the GFP were counted manually on maximum projections.

#### Movement trans-complementation assays

Cultures of *A. tumefaciens* carrying TMVΔMPΔCP-GFP plasmid highly diluted (OD_600_ 0.0001) were agroinfiltrated in *N. benthamiana* leaves together with 35S:MP-mCherry, 35S:T75A-mCherry, 35S:T75D-mCherry or free mCherry at OD_600_ 0.3. Fluorescent infection loci were observed at 7 dpi and imaged using a compact camera and UV lamp.

#### Microparticle bombardment

To study MP-GFP intercellular transport in Arabidopsis Col-0, *rda2-3* and *rda2-6* genotypes, DNA-coated gold particles were introduced in epidermal cells using microparticle bombardment. A suspension of 60 mg gold microcarriers (1.0 μm) (Bio-Rad Laboratories, Hercules, CA, USA) in 1 mL of 50% glycerol. Aliquots (35 μL) of suspended gold particles were coated with2-2.5 μg of plasmid DNA (pGGZ-35S:MP-GFP or pGGZ-35S:2xmEGFP) in the presence of 0.1 M spermidine and 2.5 M CaCl2 and vortexed for 1 min. Coated gold particles were centrifuged at 1,500 x g for 30 sec and washed twice with 70% ethanol. 35 μL of 100 % ethanol was added and vortexed until the pellet completely dispersed. This gold suspension (10 μL) was transferred onto a macrocarrier (Bio-Rad Laboratories, Hercules, CA, USA). Bombardment was performed using the PDS-1000/He Biolistic Particle Delivery System (Bio-Rad, #1652257) as described ^70^. To visualize bombardments, GFP fluorescence was monitored by confocal microscopy with excitation at 488 nm and emission detection at 495-548 nm. Maximum projection images were analyzed and cells showing GFP fluorescence were manually counted.

## Data availability

Source data have been assigned a DOI (http://doi.org/10.5281/zenodo.18468620) and are accessible via https://git.nfdi4plants.org/projects/3651. Mass spectrometry data were deposited to ProteomeXchange Consortium via the PRIDE partner repository with identifier PXD073470.

## Acknowledgements

We would like to thank Miguel A. Aranda and Eduardo Méndez-López for reading and commenting on the article. This work was supported by Ministerio de Ciencia, Innovación y Universidades (Grant PID2023-149845OA-I00) and the European Research Council (ERC) under the European Union’s Horizon 2020 research and innovation program (Grant agreement ‘SymPore’ No. 951292 to WBF and WXS) and the Alexander von Humboldt Professorship to WBF. ARE was supported in part by funds from the Interdisciplinary Graduate and Research Academy (iGRAD) Düsseldorf (Deutsche Forschungsgemeinschaft (DFG, German Research Foundation), DFG, project ID 391465903/GRK 2466).

## Author contributions

MM conceived and supervised the study. JMS and MM performed imaging and data analysis. AC and WS performed proteomics and data analysis. JMS, MM, ARE and TMS performed transport assays. MM and WBF wrote the manuscript. All authors reviewed and approved its final version.

