## Supplemental Data for "A receptor-like kinase controls plasmodesmal transport of conserved 30K viral movement proteins through phosphorylation"

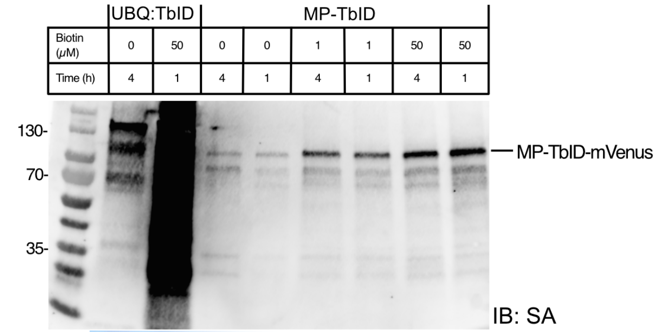


**Supplementary Figure 1. Optimization of TurboID biotinylation conditions in Arabidopsis.** Streptavidin-HRP blot showing biotinylated proteins in Arabidopsis UBQ:MP-TbID-mVenus after incubation with increasing biotin concentrations (0, 1, 50 μM) for 1 and 4 hours. UBQ:TbID plants [1] were used as a control.

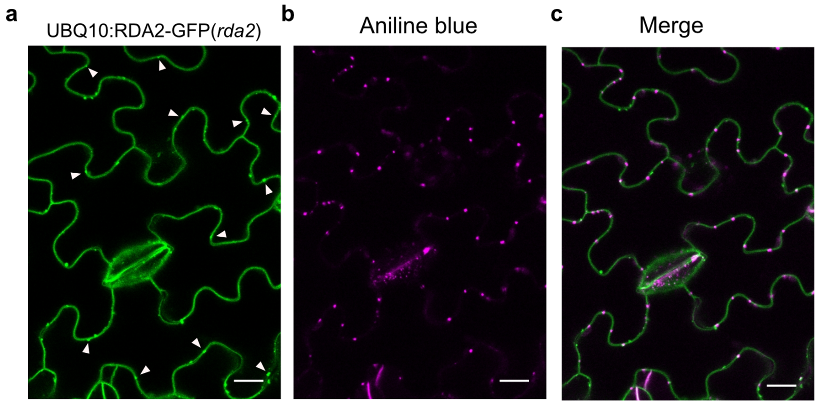


**Supplementary Figure 2. Subcellular localization of RDA2-GFP in Arabidopsis *rda2-3* complemented lines. a** Confocal image of RDA2-GFP expressed under the UBQ10 promoter in *Arabidopsis thaliana rda2-3* mutant plants. RDA2-GFP localizes predominantly to the plasma membrane and punctate structures at the cell periphery. White arrowheads indicate overlay with aniline blue. **b** Staining of plasmodesmata-associated callose using aniline blue. **c** Merged images of RDA2-GFP and aniline blue fluorescence. Scale bars: 10 µm.

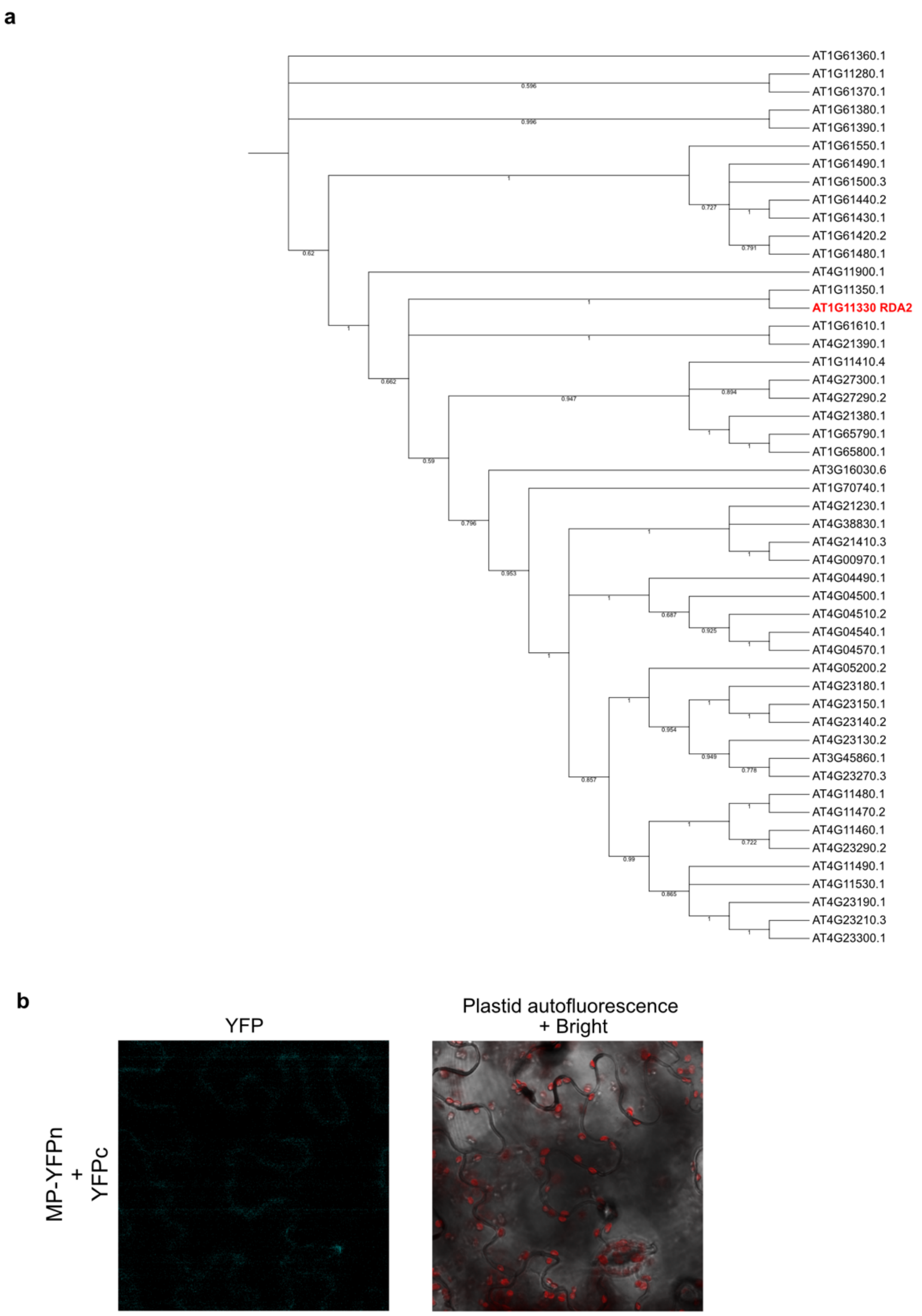


**Supplementary Figure 3. a Phylogenetic analysis of Arabidopsis RDA2 and its closest paralogs.** Phylogenetic tree of Arabidopsis RDA2 (AT1G11330, in red) and its predicted paralogs, generated with the maximum likelihood method implemented in MEGA [2]. Percent support values from 1,000 bootstrap samples are shown. The analysis identifies AT1G11350 as the closest paralog to RDA2, which was used as a negative control in the split-FP interaction assays with TMV MP. b Split YFP assay co-expressing MP-YFPn and YFPc alone, as negative control of the interaction.

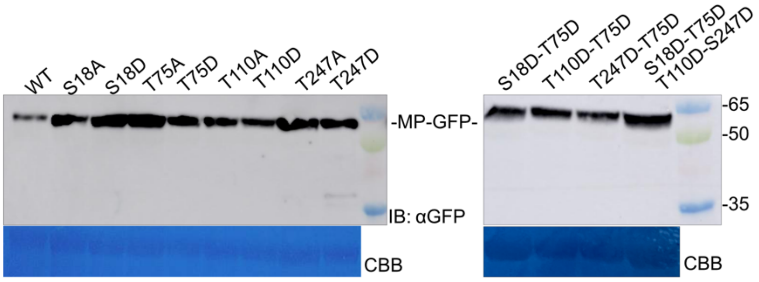


**Supplementary Figure 4. Expression of TMV MP-GFP phospho-mutants in *N. benthamiana*.** Western blot analysis of MP-GFP wild-type and phospho-mutant variants transiently expressed in *N. benthamiana*. Proteins were detected using an anti-GFP antibody to verify stability of fusion proteins. The lower panel shows the corresponding Coomassie Brilliant Blue (CBB)-stained membrane as a loading control.

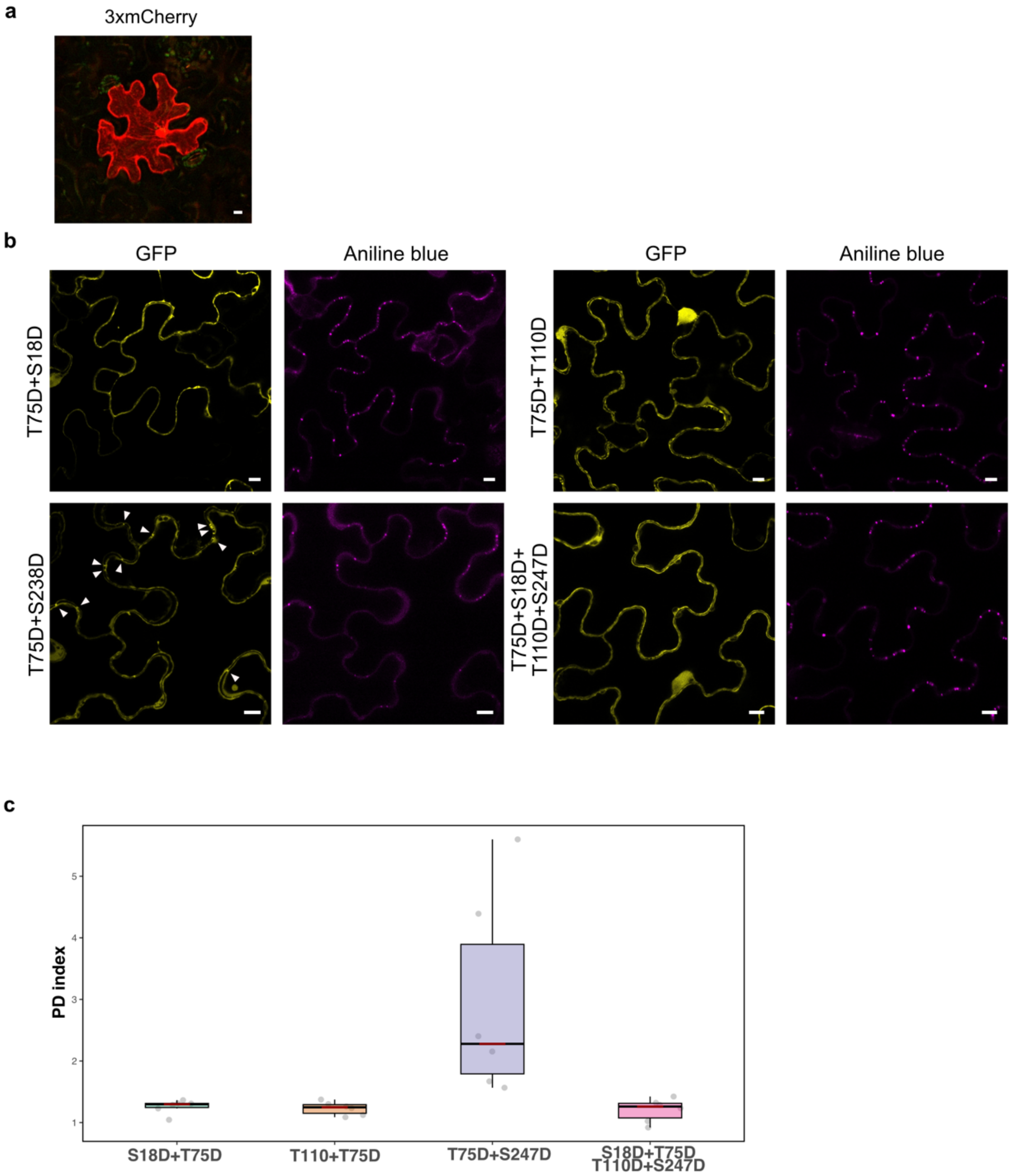


**Supplementary Figure 5. Subcellular localization of combinatorial phospho-mutants of TMV MP in *N. benthamiana*. a** Intercellular transport assay in *N. benthamiana* leaves transiently expressing a 3×mCherry tandem, which is immobile when expressed alone. **b** GFP-tagged TMV MP double mutants (S18D+T75D, T75D+T110D, T75D+S247D) and the quadruple phospho-mimetic variant were transiently expressed in *N. benthamiana* epidermal cells. Most double mutants and the quadruple mutant displayed reduced plasmodesmatal targeting and accumulated predominantly in the cytoplasm and/or nucleus. In contrast, the T75D+S247D mutant retained punctate localization at plasmodesmata, similar to the single T75D variant. Arrowheads indicate PD-associated puncta. Scale bars: 10 µm. **c** PD index of combinatorial mutants showing that T75D+S247D mutant is enriched at PD, whereas the remaining mutants display significantly lower PD association.

**Supplementary Table S1. LC-MS/MS analysis of TMV MP candidate proximal proteins**

| AGI | MAPMAN | DESCRIPTION | SUBA4 |
|---|---|---|---|
| **AJ509081** | | **MP** | |
| AT4G02450 | glycine rich proteins | glycine-rich protein | cytosol |
| AT5G35410 | signalling.calcium.CBL interacting kinase/SnRK3 | Symbols: SOS2, SnRK3.11, CIPK24 SOS2 (SALT OVERLY SENSITIVE 2) | cytosol |
| AT1G12270 | stress | stress-inducible protein, putative | nucleus |
| AT5G02890 | not assigned.no ontology | transferase family | cytosol |
| **AT1G11330** | **signalling.receptor kinases.S-locus glycoprotein like** | **S-locus lectin protein kinase family protein** | **plasma membrane** |
| AT5G46800 | transport.metabolite transporters at the mitochondrial membrane | Symbols: BOU \| BOU (A BOUT DE SOUFFLE); binding / transporter | mitochondrion |
| AT3G24830 | protein.synthesis.ribosomal protein.eukaryotic.60S subunit.L13A | 60S ribosomal protein L13A (RPL13aB | cytosol |
| AT5G23740 | protein.synthesis.ribosomal protein.eukaryotic.40S subunit.S11 | Symbols: RPS11-BETA \| RPS11-BETA (RIBOSOMAL PROTEIN S11-BETA) | cytosol |

**Supplementary Table S2. Identification of phosphorylated residues of TMV MP by LC-MS/MS**

| Position | Score | SPI | Intensity | Sequence | Modifications | m/z |
|---|---|---|---|---|---|---|
| T110 | 16.44 | 92.8 | 2.15E+04 | ADEATLGSYYtAAAK | t:Phosphorylated T | 806.3465 |
| S18 | 11.55 | 88 | 6.26E+03 | VNINEFIDLsK | s:Phosphorylated S | 693.3358 |
| S247 | 11.95 | 81.7 | 1.20E+04 | DFGGMsFKK | s:Phosphorylated S | 548.7283 |
| | 9 | 76.8 | 1.68E+04 | DFGGMsFK | | 484.6808 |
| | 8.78 | 84.4 | 1.03E+04 | DFGGMsFKK | | 366.155 |
| T75 | 4.5 | 100 | 7.62E+03 | LAGLVVtG | t:Phosphorylated T | 405.2228 |
| | 4.5 | 100 | 1.07E+04 | LAGLVVtG | | 405.2231 |
| | 4.5 | 100 | 5.82E+03 | LAGLVVtG | | 405.2238 |

Note: the identified phosphorylation site was marked in red

**Supplementary Table S3. Identification of phosphorylated residues of CMV MP by LC-MS/MS**

| Position | Score | SPI | Intensity | Sequence | Modifications | m/z |
|---|---|---|---|---|---|---|
| S128 | 7.74 | 66.6 | 1.39E+03 | GSLRIYLADLGDKELSPIDGQCVsLHNHDLPALVSFQP | s:Phosphorylated S | 1050.2419 |
| | 6.26 | 63.9 | 1.43E+04 | GSLRIYLADLGDKELSPIDGQCVsLHNHDLPALVSFQP | | 1050.2419 |
| | 5.95 | 69.7 | 1.39E+03 | GSLRIYLADLGDKELSPIDGQCVsLHNHDLPALVSFQP | | 1050.2421 |
| S106 Y110 | 6.69 | 67.9 | 2.52E+03 | RTVSTDAEGsLRIyLAD | s:Phosphorylated S y:Phosphorylated Y | 487.49 |
| | 6.09 | 67.8 | 2.86E+03 | RTVSTDAEGsLRIyLADL | | 535.7646 |
| | 5.99 | 79.1 | 1.01E+04 | RTVSTDAEGsLRIyLADL | | 535.7646 |
| S73 Y75 | 6.31 | 65.9 | 9.44E+02 | FKsGyDVGELCSKGYMSVP | s:Phosphorylated S y:Phosphorylated Y | 748.3123 |

Note: the identified phosphorylation site was marked in red

**Supplementary Table S4. Primers used in this study**

| Name | Sequence (5´-3´) | Description |
|---|---|---|
| TMV_MP_Fw | TTTGGTCTCAGGCTCTATGGCTCTAGTTGTTAAAGG | For cloning TMV MP in pGGC |
| TMV_MP_Rv | TTTGGTCTCACTGAAAGTCATTAAAACGAATCCGATTCGG | |
| RDA2_ Fw | TTTGGTCTCAGGCTCTATGGTGGTTTCAGTGACCA | For cloning RDA2 in pGGC |
| RDA2_ Rv | TTTGGTCTCACTGAACGTCCTGTTACAGCTGTGAG | |
| RDA2_TM_KIN_Fw | TTTGGTCTCAGGCTCTATGGCTCATTCAGAACTCAAAACAC | For Split-FP |
| RDA2_KIN_Fw | TTTGGTCTCAGGCTCTATGCCAGCTCCAGCGAAAG | |
| RDA2_deltaTM_Rv | TTTGGTCTCACTGACTGCTAGATTGCTATGTGTTTT | |
| AT1G11350.1_Fw | TTTGGTCTCAGGCTCTATGGGTTGTTTGCTAATTCTTC | |
| AT1G11350.1_Rv | TTT GGTCTC A CTGAACGTCCTGTTATTTTTGTGAG | |
| TMV_qPCR_Fw | AAGTGTCTCCCAGGCAACAG | For qPCR for detection of viral RNA, specific for RdRp |
| TMV_qPCR_Rv | TTAGGCGCAGCAACAGACTT | |
| Actin2 1093 Fw | GTGGTCGTACAACCGGTATT | |
| Actin2 1288 Rv | CACGTCCAGCAAGGTCAAGA | |
| MP_CMV-fny_Fw | TTTGGTCTCAGGCTCATGGCTTTCCAAGGTACC | For cloning CMV MP in pGGC |
| MP_CMV-fny_Rv | TTTGGTCTCACTGAAAGACCGTTAACCACCTG | |
| TMV_mimet247_Fw | GATTTTGGAGGAATGGATTTTAAAAAGAATAAT | For TMV MP mutants |
| TMV_mimet247_Rv | ATTATTCTTTTTAAAATCCATTCCTCCAAAATC | |
| TMV_no-mimet247_Fw | GATTTTGGAGGAATGGCTTTTAAAAAGAATAAT | |
| TMV_no-mimet247_Rv | ATTATTCTTTTTAAAAGCCATTCCTCCAAAATC | |
| TMV_mimet75_Fw | GCCGGTTTGGTCGTCGATGGCGAGTGGAACTTG | |
| TMV_mimet75_Rv | CAAGTTCCACTCGCCATCGACGACCAAACCGGC | |
| TMV_no_mimet75_Fw | GCCGGTTTGGTCGTCGCTGGCGAGTGGAACTTG | |
| TMV_no-mimet75_Rv | CAAGTTCCACTCGCCAGCGACGACCAAACCGGC | |
| TMV_no_mimet_T110_Fw | CTCGGATCTTACTACGCTGCAGCTGCAAAGAAA | |
| TMV_no_mimet_T110_Rv | TTTCTTTGCAGCTGCAGCGTAGTAAGATCCGAG | |
| TMV_no_mimet_S18_Fw | GAGTTTATCGACCTGGCTAAAATGGAGAATATC | |
| TMV_no_mimet_S18_Rv | GATATTCTCCATTTTAGCCAGGTCGATAAACTC | |
| TMV_mimet_T110_Fw | CTCGGATCTTACTACGATGCAGCTGCAAAGAAA | |
| TMV_mimet_T110_Rv | TTTCTTTGCAGCTGCATCGTAGTAAGATCCGAG | |
| TMV_mimet_S18_Fw | GAGTTTATCGACCTGGATAAAATGGAGAATATC | |
| TMV_mimet_S18_Rv | GATATTCTCCATTTTATCCAGGTCGATAAACTC | |

| | | |
|---|---|---|
| Replicon_TMV_Fw | GGTTACCTAAATAATAGACGGAGGGCCCATGGAAC | For TMVΔMPΔCP-GFP replicon |
| Replicon_TMV_Rv | TCTATTATTTAGGTAACCTTTGTCAGGTCGATAAACTC | |
| 2xmEGFP_Fw | GGGGACCACTTTGTACAAAAAGCAGGCTTCACCATGGTTTCTAAGGGTGAGG | For microparticle bombardment |
| 2xmEGFP_Rv | GGGGACCACTTTGTACAAGAAAGCTGGGTGTTACTTGTACAGCTCGTCCATGCC | |
